# The convoluted evolutionary history of the capped-golden langur lineage (Cercopithecidae: Colobinae) – concatenation versus coalescent analyses

**DOI:** 10.1101/508929

**Authors:** Kunal Arekar, Abhijna Parigi, K. Praveen Karanth

**Author notes:** Corresponding author. Email –.

## Abstract

Evolutionary studies have traditionally relied on concatenation based methods to reconstruct relationships from multiple markers. However, due to limitations of concatenation analyses, recent studies have proposed coalescent based methods to address evolutionary questions. Results from these methods tend to diverge from each other under situations where there is incomplete lineage sorting or hybridization. Here we used concatenation as well as multispecies coalescent (MSC) methods to understand the evolutionary origin of capped and golden langur (CG) lineage. Previous molecular studies have retrieved conflicting phylogenies, with mitochondrial tree grouping CG lineage with a largely Indian genus *Semnopithecus,* while nuclear markers support their affinities with a Southeast Asian genus, *Trachypithecus*. However, as pointed by others, the use of nuclear copies of mitochondrial DNA in the above studies might have generated the discordance. Because of this discordance, the phylogenetic position of CG lineage has been much debated in recent times. In this study, we have used nine nuclear and eight mitochondrial markers. Concatenated nuclear as well as the mitochondrial dataset recovered congruent relationships where CG lineage was sister to *Trachypithecus*. However nuclear species tree estimated using different MSC methods were incongruent with the above result, suggesting presence of incomplete lineage sorting (ILS)/hybridisation. Furthermore, CG lineage is morphologically intermediate between *Semnopithecus* and *Trachypithecus*. Based on this evidence, we argue that CG lineage evolved through hybridisation between *Semnopithecus* and *Trachypithecus*. Finally, we reason that both concatenation as well as coalescent methods should be used in conjunction for better understanding of various evolutionary hypotheses.

## 1. Introduction

This day and age most phylogenetic studies use multiple markers to reconstruct the evolutionary relationships between taxa. Typically, such studies employ concatenated analysis which involves building a supermatrix by combining multilocus molecular data (de Querioz & Gatesy, 2006). However, the basic assumption of this approach, which is that all the loci experience same the evolutionary history, is often incorrect (de Querioz & Gatesy, 2006). Processes such as hybridisation and incomplete lineage sorting (ILS) can lead to discrepancy between the gene trees and the true species tree (Degnan James H., Rosenberg, 2006; Maddison, 1997). To overcome this limitation of the concatenation analysis, coalescence-based tree building methods have been proposed as an alternative (Degnan & Rosenberg, 2009; Knowles, L.L, Kubatko, 2010; Song, Liu, Edwards, & Wu, 2012; Zhong, Liu, & Penny, 2014).

ILS arises when two or more gene trees fail to coalesce back in time in the most recent common ancestor (Maddison, 1997) and this is more likely to occur in lineages where not enough time has passed since speciation (Leache & Rannala, 2011) whereas, hybridisation is a process that results in mixing of previously isolated gene pools. The interbreeding of individuals from different gene pools results in a hybrid species that shares genetic information with both the parent species (Mallet, 2005, 2007; Zinner, Arnold, & Roos, 2011). There have been many methods developed to estimate species tree in presence of ILS (Bryant, Bouckaert, Felsenstein, Rosenberg, & Roychoudhury, 2012; Chifman & Kubatko, 2015; Edwards, Liu, & Pearl, 2007; J. Heled & Drummond, 2010; Laura S. Kubatko, Carstens, & Knowles, 2009; Liu & Pearl, 2007; Liu, Yu, & Edwards, 2010; Mirarab & Warnow, 2015; Mossel & Roch, 2008) and hybridisation (Gauthier & Lapointe, 2007; Jin, Nakhleh, Snir, & Tuller, 2006; Olave, Avila, Sites, & Morando, 2017; Posada, 2002; Xu, 2000). However, distinguishing between these two processes is difficult and very few methods have been developed that incorporates both hybridisation and ILS (Buckley, Cordeiro, Marshall, & Simon, 2006; Joly, McLenachan, & Lockhart, 2009; Meng & Kubatko, 2009; Than, Ruths, Innan, & Nakhleh, 2007). Importantly, multispecies coalescent (MSC) models incorporate ILS in species tree estimation but these methods cannot distinguish between hybridisation and ILS (L. S. Kubatko, 2009). Additionally, there is a dearth of studies that compare and contrast concatenated analysis vs. MSC analysis under conditions where ILS/hybridisation is suspected.

Here we have used multiple nuclear and mitochondrial markers in conjunction with different tree building approaches to better understand the processes that gave rise to a lineage of Asian colobine monkeys – the capped and golden langur (CG) lineage. Colobines are predominantly leaf eating monkeys (Chivers & Hladik, 1980) distributed in the tropical old world. They are a diverse group consisting of 10 genera and over 50 species (Brandon-Jones et al., 2004; Groves, 2001). Among the Asian colobines, *Semnopithecus* and *Trachypithecus*, are the most species rich genera distributed in the Indian subcontinent and Southeast Asia (SEA) respectively. There has been much ambiguity in the taxonomy of these two genera (Brandon-Jones et al., 2004; Davies & Oates, 1994; Groves, 2001); however, with the advent of molecular tools many of these issues have been resolved(Ashalakshmi, Nag, & Karanth, 2014; Karanth, 2008; Karanth, Singh, Collura, & Stewart, 2008). Nevertheless, the taxonomic placement of two species in the genus *Trachypithecus* continues to be in debate; these include the endangered golden langur (*Trachypithecus geei*) and capped langur (*Trachypithecus pileatus*). Golden langur, capped langur and Shortridge’s langur (*Trachypithecus shortridgei*) are the three species that are together called as *T. pileatus* group(Wang et al., 2015) However, in this study, we have included *Trachypithecus shortridgei* within CG lineage; recently, *T*. *shortridgei* was split from capped langur and elevated to species level(Brandon-Jones et al., 2004; Groves, 2001).

Golden langur is distributed in parts of Bhutan and adjoining Indian state of Assam in Northeast India(Ram et al., 2016; Wangchuk, Inouye, & Hare, 2008) whereas, capped langur is more widespread and is distributed in northeast India, parts of Bhutan, Bangladesh, northwest Myanmar and southern China (Kumar & Solanki, 2008; Srivastava & Mohnot, 2001; Zhang, Wang, & Quan, 1981). Molecular data suggests the two species are sister lineages, however their phylogenetic position in relation to *Semnopithecus* and *Trachypithecus* remains unresolved. In the molecular studies undertaken by two separate groups, the mitochondrial Cytochrome *b* (Cyt-*b*) phylogenetic tree placed Capped-Golden (CG) lineage in the *Semnopithecus* clade whereas nuclear autosomal and Y chromosomal markers suggested that they were sister to *Trachypithecus* (Karanth et al., 2008; Osterholz, Walter, & Roos, 2008). This discordance led the authors to hypothesise reticulate evolution of CG lineage due to past hybridisation between *Semnopithecus* and *Trachypithecus*. Osterholz et al. (2008) pointed out that incomplete lineage sorting in either nuclear or mitochondrial markers could also generate this discordance. Interestingly the distribution of CG lineage is flanked by *Semnopithecus* on the west and by *Trachypithecus* in the east (Groves 2001; Karanth et al., 2008; Osterholz et al., 2008). The same year another study looked at the phylogenetic position of CG lineage using Cyt-*b* marker (Wangchuk et al., 2008). Surprisingly in their mitochondrial tree CG lineage branched within the *Trachypithecus* clade. Thus, Wangchuk et al. (2008) results were consistent with the retrieved topology using nuclear markers (Karanth et al., 2008; Osterholz et al., 2008) of the aforementioned studies and did not support the mitochondrial-nuclear discordance scenario (Karanth et al., 2008; Osterholz et al., 2008). Accordingly, they rejected the hybridisation hypothesis as a possible reason for the origin of CG lineage.

These studies suggested that CG lineage had at least two types of Cyt-*b* sequences, one of which cluster them with *Semnopithecus* and the other with *Trachypithecus*. Multiple mitochondrial sequences when amplified from the same species suggest the presence of nuclear mitochondrial DNA sequences (numts). Numts are mitochondrial sequences incorporated into the nuclear genome(Bensasson, Zhang, Hartl, & Hewitt, 2001; Blanchard & Lynch, 2000; Ford Doolittle, 1998) and behave like nuclear pseudogenes. These nuclear copies of mitochondrial sequences when considered as true mitochondrial sequences and used in analysis can lead to erroneous phylogenetic topologies (Bensasson et al., 2001; Collura & Stewart, 1995). Numts have been reported from both *Semnopithecus* and *Trachypithecus* (Karanth, 2008). A recent study reported the isolation of full length numts from capped langur and *Trachypithecus shortridgei* (Wang et al., 2015). These numt sequences branched with the genus *Semnopithecus* (Karanth et al., 2008; Osterholz et al., 2008), but the mitochondrial sequences from capped langur and *Trachypithecus shortridgei* places them sister to other members of *Trachypithecus*, consistent with the Cyt-*b* phylogeny of Wangchuk et al., (2008). Thus, the authors concluded that “mitochondrial” Cyt-*b* sequences of CG lineage used by Karanth et al. (2008) and Osterholz et al. (2008) were numts.

Taken together, these studies would suggest that CG lineage indeed belongs to *Trachypithecus* clade. Nevertheless, studies on numts by Karanth (2008) and Wang et al. (2015) suggest that hybridisation hypothesis for the origin of CG lineage cannot be rejected. Karanth (2008) isolated numt sequences from capped langur, golden langur and Phayre’s leaf monkey (*T. phayrei*). Assuming that CG lineage belongs to *Trachypithecus*, these numts are predicted to be related to the *Trachypithecus* clade or expected to be sister to the clade consisting of both *Semnopithecus* and *Trachypithecus*. Interestingly in their analysis some numts isolated from both capped and golden langurs were also sister to *Semnopithecus*. This unusual placement of some CG numts thus suggested hybridisation. Based on molecular dating analysis Karanth (2008) estimated the hybridisation to have occurred between 7.1 to 3.4 million years ago (mya). Similarly, Wang et al. (2015) proposed that the numts originated in the genus *Semnopithecus* and hypothesised a unidirectional introgression from *Semnopithecus* into the ancestors of capped langur and *T*. *shortridgei*. They dated the time of hybridisation to be no later than 0.26 mya and no earlier than 3.47 mya. Hybridisation in primates has been reported between species and subspecies(Pastorini, Zaramody, Curtis, Nievergelt, & Mundy, 2009; Rabarivola, Meyers, & Rumpler, 1991; Vasey & Tattersall, 2002), but there are a few studies that have observed it between genera (Davenport et al., 2006; Dunbar & Dunbar, 1974; Jones et al., 2005; Zinner, Chuma, Knauf, & Roos, 2018).

Thus, nuclear and mitochondrial data support placing CG lineage in *Trachypithecus*. Whereas, the numts data suggests past hybridisation between *Trachypithecus* and *Semnopithecus*. Nevertheless, it must be noted that these studies have undertaken phylogenetic analyses of concatenated markers to ascertain the phylogenetic position of CG lineage. Such analyses, as mentioned above, are inappropriate when markers have disparate evolutionary histories. Therefore, here we compare the phylogenetic position of CG lineage determined using both concatenated phylogenetic analyses and species tree approaches (MSC analyses) based on nine nuclear markers to better understand the evolution of this enigmatic taxon. For the nuclear analyses, multiple samples of capped and golden langurs have been used. Furthermore, we also undertook a separate analysis of mtDNA dataset which included a large mitochondrial fragment from a wild caught capped langur to reconfirm its position in the mitochondrial tree. This was important given the past studies either used smaller fragment of mtDNA or included zoo derived specimen of species from CG lineage.

## 2. Materials and Methods

### 2.1. Taxon sampling and DNA extraction

Two fecal samples from golden langur and four samples (two fecal and two tissue) from capped langurs were collected non-invasively from different zoos across India (Table S4). Fresh faeces were collected only from wild caught captives, using plant twigs to avoid contamination and preserved in absolute alcohol (Merck, Germany) and stored in −20°C.

For DNA extraction from tissue samples, we used the commercially available DNeasy® Blood & Tissue Kit (QIAGEN Inc.), we followed the manufacturer’s protocol. DNA from fecal samples was extracted using the commercially available QIAamp DNA stool mini kit (QIAGEN Inc.), following the manufacturer’s protocol with slight modifications as mentioned in Mondol et al. (2009), however, we did not add the carrier RNA (Poly A)(Kishore, Reef Hardy, Anderson, Sanchez, & Buoncristiani, 2006; Mondol et al., 2009). Each extraction had a negative control to monitor contamination. The quantity of extracted DNA was measured using a NanoDrop 2000 Spectrophotometer (Thermo Fisher Scientific Inc).

### 2.2. PCR Amplification and Sequencing

Nine nuclear markers were selected for this study (Table 1) (Perelman et al., 2011; Karanth et al., 2008). The nuclear markers were amplified using the following conditions. A 25 µl PCR reaction volume was set with 2 µl (total DNA concentration varied between samples ranging from 20 ng/µl to 80 ng/µl) of DNA extract, 0.25 mM of dNTPs (Bangalore Genei, Bangalore), 0.2 µM of each primer (Amnion Biosciences, Bangalore), 1U Taq polymerase (New England BioLabs® Inc.) and a standard 1X reaction buffer premixed with 1.5 mM MgCl_2_ (New England BioLabs® Inc.). A ‘touchdown’ PCR program was used with an initial annealing temperature of 60 °C with subsequent decrease by 1 °C per cycle till the specified annealing temperature, 50 °C, is reached. Each reaction was performed for 95 °C for 15 sec, 60-50 °C for 30 sec and 72 °C for 60 sec, with initial denaturation at 95 °C for 10 min. PCR products were purified using the QIAquick® PCR purification kit (QIAGEN Inc.) following the manufacturer’s protocol, for some samples we used ExoSAP-IT (Affymetrix®, USB Products). The purified product was outsourced for sequencing to Amnion Biosciences Pvt. Ltd., Bangalore; some of the samples were outsourced to Medauxin, Bangalore.

**Table 1:**
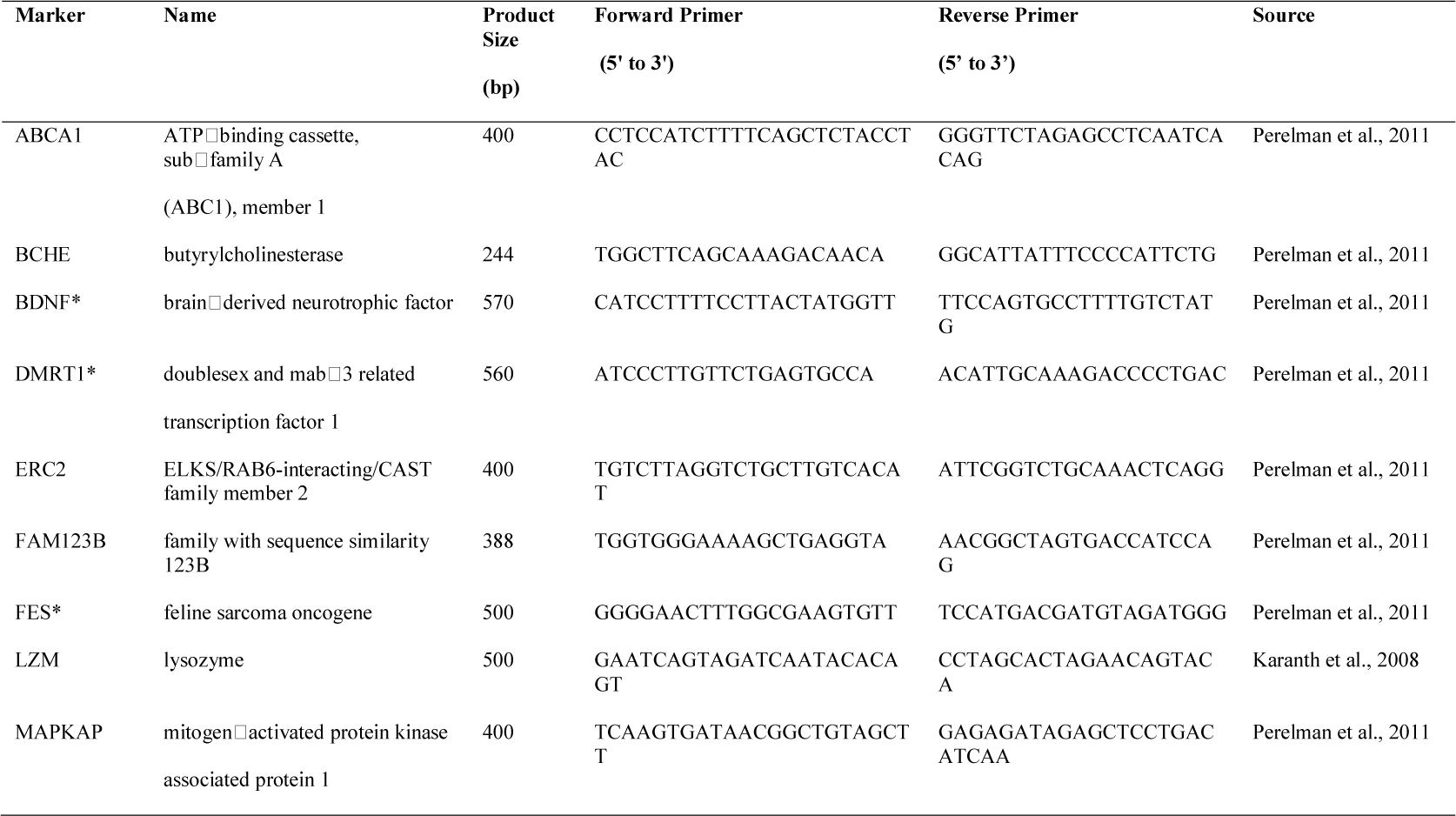
Nuclear markers used in this study with primers and source. * indicates primer pairs that have not been redesigned.

Additionally, eight mitochondrial markers were amplified from a single capped langur tissue sample (Table 2). First, we used a long-range PCR kit (Expand Long Template PCR System, Roche) to amplify a long fragment of mitochondrial DNA (mtDNA) genome in order to minimise the chances of amplifying a numt sequence. Second, we identified these eight genes by comparing our sequence with the whole mitochondrial (mt) genome of capped langur available in GenBank. Following primer pair was used to amplify the long fragment; 5275F (5′ – ACCYCTGTCTTTAGGTTTACAGCCTAATG – 3′) and 11718R (5′ – CCAATGGATAGCTGTTATCCTTTAAAAGTTGAG – 3′) (Raaum, Sterner, Noviello, Stewart, & Disotell, 2005). The PCR cycle conditions followed were as per the manufacturer’s protocol (Roche). PCR product was outsourced for purification and sequencing to Amnion Biosciences Pvt. Ltd., Bangalore. We were not able to amplify the long mitochondrial fragment from fecal samples because of the low yield of degraded DNA which is typical of faecal samples. We did not attempt to amplify small mitochondrial fragments for single markers from the fecal samples, to avoid accidental amplification of numt sequences.

**Table 2:**
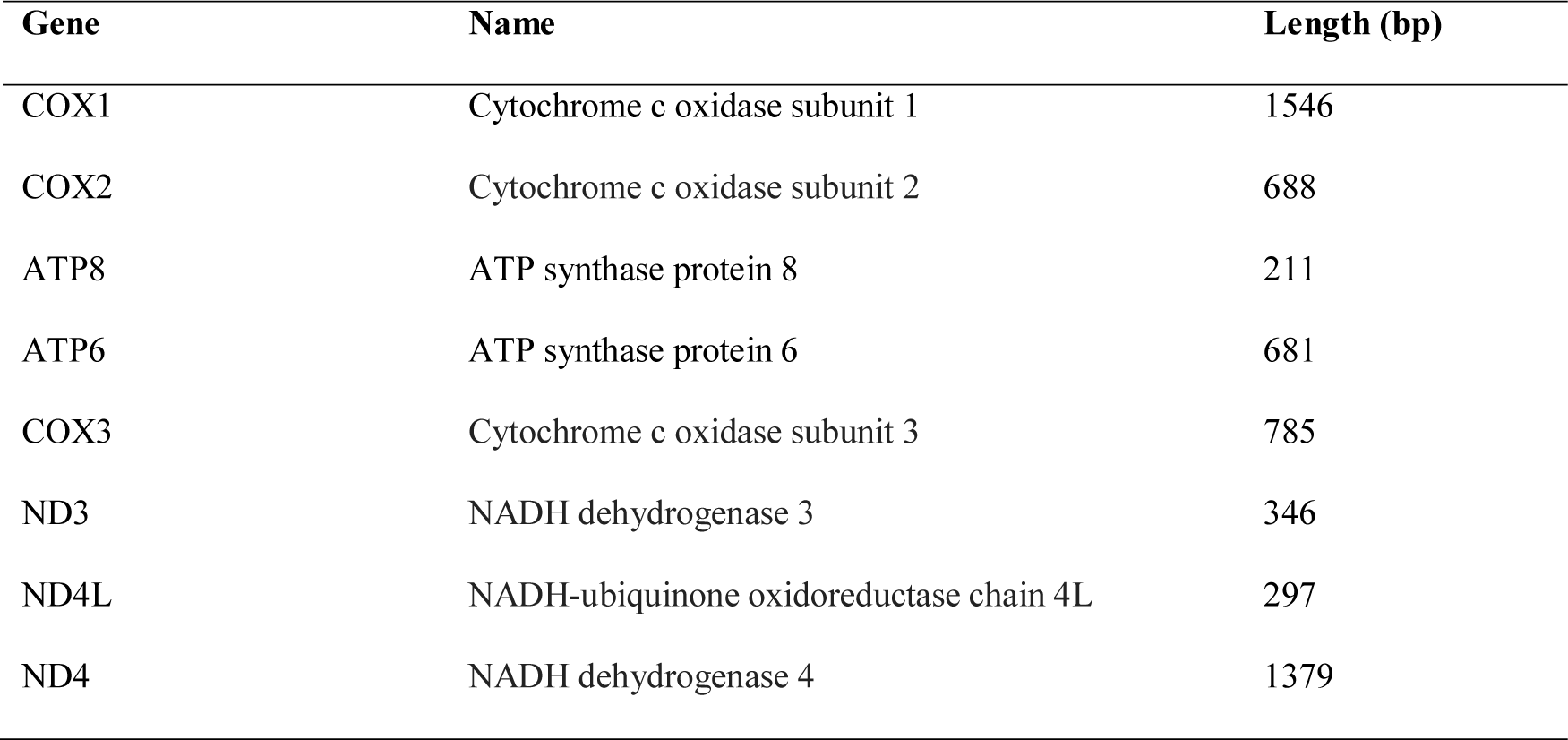
Eight genes that are amplified in the long mitochondrial DNA (mtDNA) fragment (5933 bp).

### 2.3. Phylogenetic analysis

The sequence chromatogram file was viewed and edited manually using ChromasLite v2.01 (Technelysium Pty Ltd). We generated a long fragment of mitochondrial genome for one capped langur tissue sample (CES11/299, Table S4). This long fragment of mtDNA consists of eight protein coding loci – Cox1, Cox2, ATP8, ATP6, Cox3, ND3, ND4L, ND4 (Table 2). All the sequences are deposited in GenBank (Table #; sequences will be deposited after the acceptance of manuscript). Each sequence was checked by comparing with orthologous sequences available in GenBank by using Nucleotide BLAST search tool.

The mitochondrial fragment, sequenced from one capped langur (belong to the CG lineage group) tissue sample in this study, was aligned with 19 Asian colobines and one African colobine sequences available in GenBank (Table S1). Downloaded sequences consisted of representatives from odd-nosed monkeys, *Trachypithecus* group, *Semnopithecus* group and *Colobus* used as an African colobine outgroup. Furthermore, the six non-coding tRNAs from the mitochondrial fragment were excluded from the analysis. Nuclear dataset generated in this study i.e. seven sequences from individuals that belong to CG lineage and one sequence from *Semnopithecus hypoleucos*, were aligned with sequences of 13 other Asian and African colobines available in Genbank (Table S2). The nine nuclear genes were concatenated to create a multilocus supermatrix of the nuclear data. The two data sets were aligned separately using Muscle algorithm (Edgar, 2004) incorporated in MEGA v5.2 (Tamura et al., 2011) with default settings. We used p-distance algorithm in MEGA to estimate the evolutionary divergences between sequences.

PartitionFinder 1.1.0 (Lanfear, Calcott, Ho, & Guindon, 2012) was used to pick the best partition scheme as well as the model of sequence evolution. Partitioning schemes used for analysis of our data are listed in Table S3 [(A) and (B)]. Phylogenetic reconstruction was performed using Maximum Likelihood (ML) and Bayesian methods. ML analysis was performed in RAxML 7.4.2 incorporated in raxmlGUIversion 1.3 (Stamatakis, 2006). As there is no provision in RAxML for applying multiple models across partitions, we used GTR + G as the model of substitution for all partitions in our data set (Table S3). We performed 1000 slow bootstrap replicates to assess support for different nodes. The Bayesian analysis was done in MrBayes 3.2.2 (Ronquist et al., 2012). Nuclear data analysis was run for 15 million generations with sampling frequency at 500 and the mitochondrial dataset was run for 15 million generations and sampling frequency at 2000. Analyses had two parallel runs with four chains, convergence of the runs was determined when the standard deviation of split frequencies was <0.01. Using Tracer v1.6 (Rambaut, A., Suchard, M.A., Xie, D., Drummond, 2013) we plotted each parameter against generation time to determine the effective sample size (ESS) value of >200. The first 25% trees were discarded as burn-in.

### 2.4. Species tree estimation

Evolutionary processes like hybridisation, incomplete lineage sorting (ILS), gene duplication and gene loss make it difficult to estimate species trees from multi-locus nuclear data. These processes can make gene trees incongruent and different from the overall species tree (Maddison, 1997). The individual gene trees from the nine nuclear markers were uninformative due to lack of phylogenetically informative sites; therefore, to infer a nuclear marker based species tree we used two different approaches. First the program ASTRAL II (Mirarab and Warnow, 2015) was used to build species tree for the nuclear dataset. It uses individual nuclear gene trees as input file and estimates a phylogeny that agrees with the maximum quartet trees induced by the gene tree set. The command “java -jar astral.4.10.11.jar –I in.tree -o out.tre” was used to run the input file and to get an output tree file. Input tree file was generated by combining individual gene tree outputs from RAxML.

ASTRAL is a ‘summary statistic’ method which uses input gene trees to estimate species tree. However, summary methods are known to be sensitive to gene tree estimation error, specifically on short alignments (Chou et al., 2015). The program SVDquartets is a ‘single site’ method that first builds a quartet tree for all possible quartet combinations; for all the possible trees in a quartet, SVD (single value decomposition) score is calculated and the relationship with the lowest score is picked up as the best tree. It then uses a quartet assembly technique, like QFM (Reaz, Bayzid, & Rahman, 2014) to build a tree. We ran the program with 100000 quartets to evaluate with the option to check all the possible quartets in PAUP* 4.0a157 (Swofford, 2002). Two approaches were used; first species tree analysis was undertaken with “taxa partition” option wherein individual members were assigned to major lineages recovered from the concatenation analysis (Fig. 1 & Fig. 2) and secondly, the analysis was run without taxon partitions where the tips were designated as separate populations rather than grouping them in major lineages. For both the analysis, we used the multispecies coalescent tree model and the analyses were subjected to 100 bootstrap replicates.

**Fig. 1:**
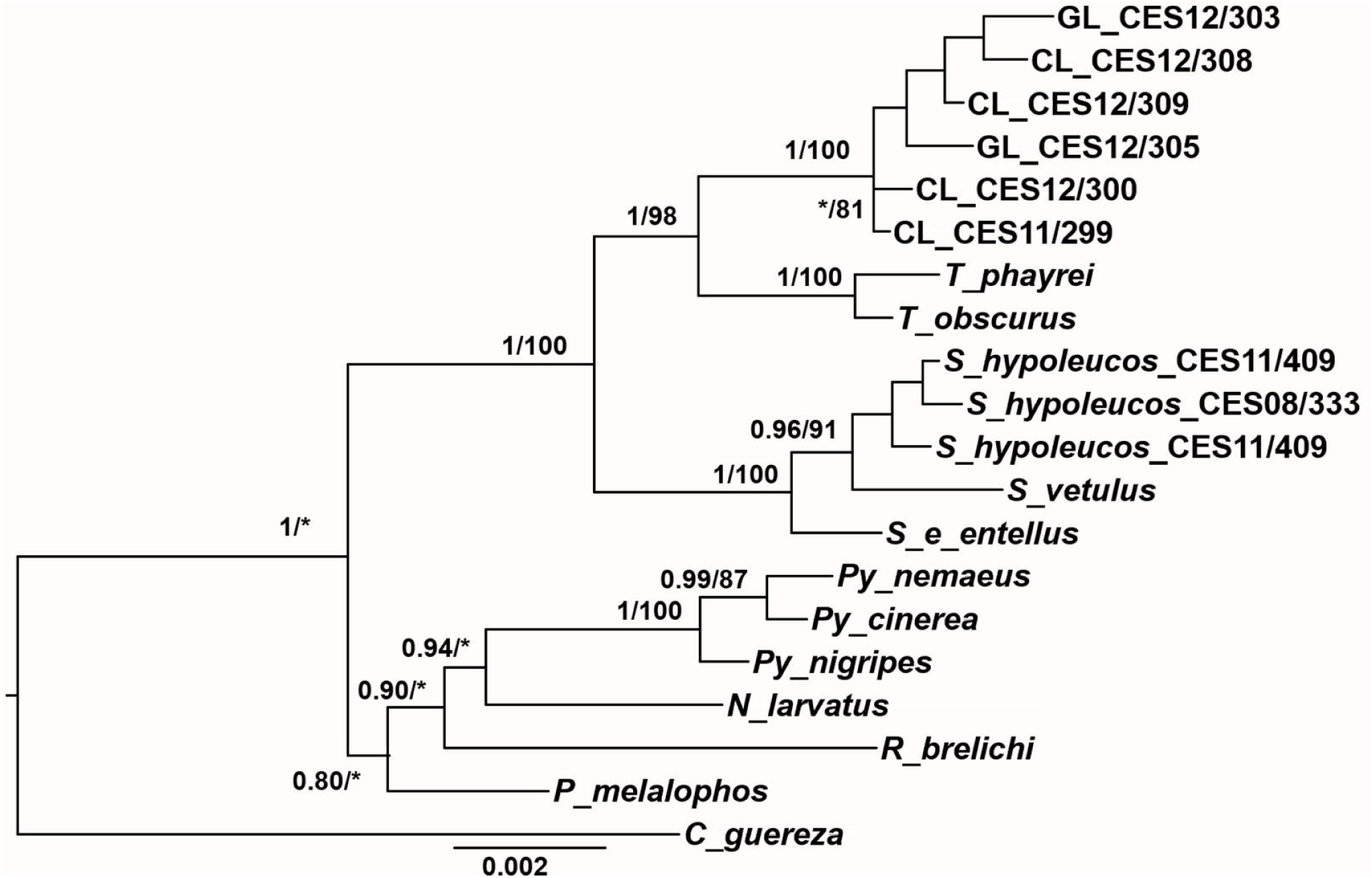
Bayesian phylogeny of Asian colobines for the concatenated nuclear data set. The numbers on the nodes are the posterior probability values/Maximum-likelihood bootstrap support. Support is not shown for those nodes which had both the Posterior probability and ML bootstrap less than 0.75 and 75, respectively. (* indicates Posterior probability or ML bootstrap values less than 0.75 or 75 respectively for that particular node only) CL = capped langur; GL = golden langur. Support for all the nodes within the CL and GL clade was below 0.75/75.

**Fig. 2:**
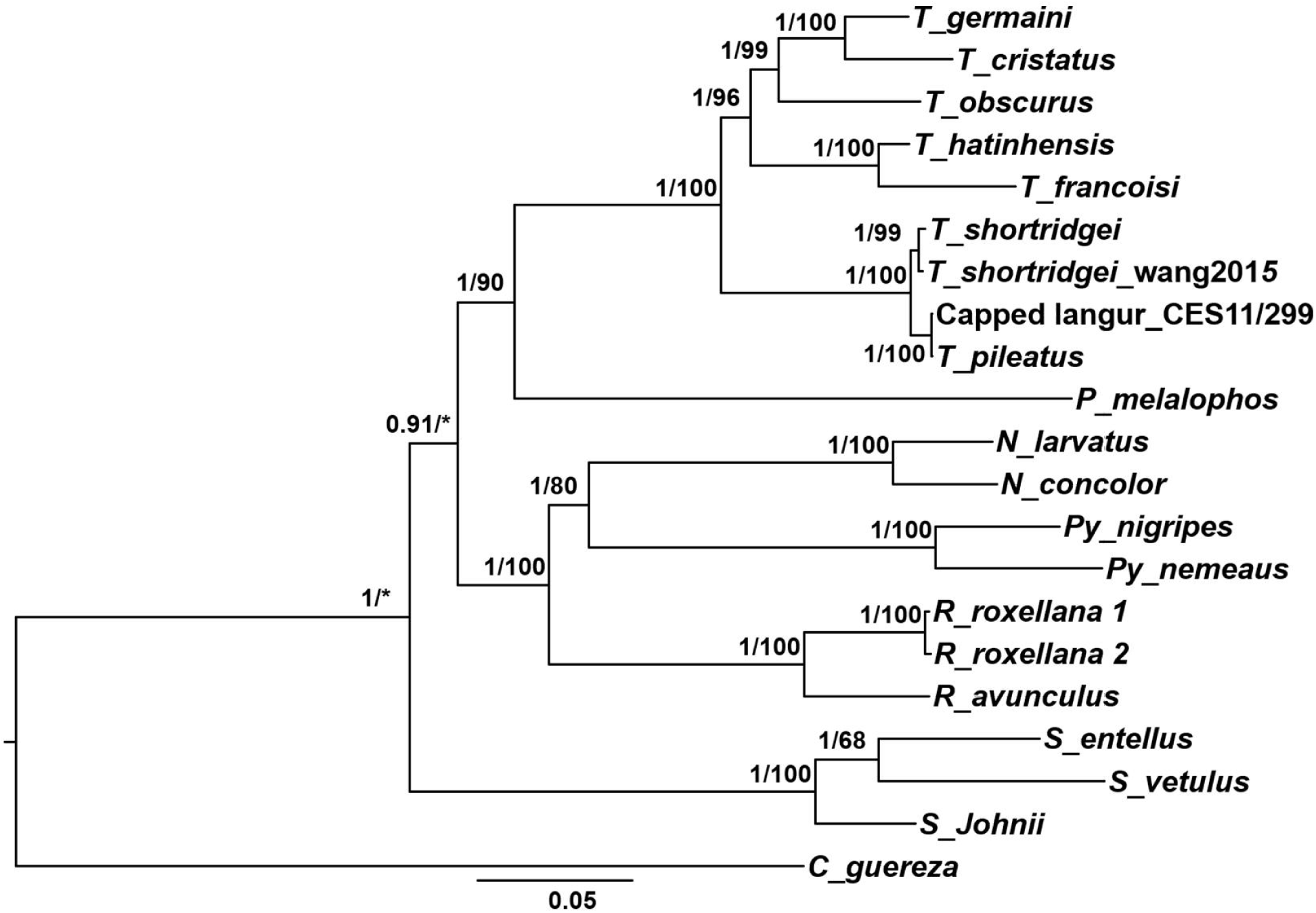
Bayesian phylogeny of Asian colobines for the mitochondrial concatenated data set. The values at the nodes denote the posterior probability values/Maximum-likelihood bootstrap support. Support is not shown for those nodes which had both the Posterior probability and ML bootstrap less than 0.75 and 75, respectively. (* indicates Posterior probability or ML bootstrap values less than 0.75 or 75 respectively for that particular node only). In this tree, the capped langur sequence was generated in this study and the *T. pileatus* was downloaded from GenBank (KF680163)

### 2.5. Divergence dating

We used the species tree ancestral reconstruction - *BEAST (Joseph Heled, Bouckaert, Drummond, & Xie, 2013) incorporated in BEAST v2.4.7 (Bouckaert et al., 2014) for divergence time estimation for our nuclear data set. Input files were created in BEAUti v2.4.7 (Bouckaert et al., 2014). The selected partitioning scheme is described in Table S3(A). We used an uncorrelated relaxed lognormal clock for each gene with default values for all the priors. Yule model was used as species tree prior. It is important to set the correct ploidy levels as differences in effective population size influences the coalescence time (Charlesworth, 2009). All the genes in our analysis are autosomal nuclear except for FAM123B, which is an X-linked locus; so, the ploidy levels were selected accordingly. We *a priori* grouped the tips into their respective genera; individuals from CG lineage were assigned to a separate group (Table S5). Two independent analyses were run for 80 million generations, sampling every 5000 generations. Stationarity was assessed in Tracer v1.6 (Rambaut et al., 2013) based on the plots of parameters vs. state or generation and the effective sample sizes (>200). The tree files were combined using LogCombiner 2.4.7 and constructed maximum clade credibility tree with median height using TreeAnnotator v2.4.7 after discarding first 25% trees as burn-in.

We used two fossil calibrations and one secondary calibration for dating the phylogeny. A previous study (Perelman et al., 2011) inferred the most recent common ancestor (MRCA) for Asian colobine at 8.8mya with 95% credible interval (CI) of 6.5 – 11.2 mya. Therefore, the molecular clock prior was set to normal with mean of 8.8. One of the fossils we used for calibration is a primate from middle to late Miocene Eurasia, *Mesopithecus pentelicus* (Pan, Groves, & Oxnard, 2004). To the best of our knowledge, it has never been used as a calibration for any primate phylogeny. According to Pan et al. (2004), this fossil is related to the odd-nosed colobine genus *Rhinopithecus* and *Nasalis*. According to our ML and Bayesian phylogenies, *Pygathrix* is placed sister to the *Rhinopithecus* + *Nasalis* clade, hence we constrained the *Rhinopithecus* + *Nasalis* + *Pygathrix* clade as monophyletic and calibrated this node using lognormal prior distribution with mean = 9.0, offset = 6.0. Another fossil we used is a late Pliocene *Semnopithecus gwebinensis* fossil discovered from the Irrawaddy sediments of central Myanmar (Takai et al., 2016). For this calibration, we constrained *Semnopithecus* + *Trachypithecus* + CG lineage as monophyletic and used a lognormal distribution prior with mean = 12.0 and offset = 2.6.

## 3. Results

### 3.1. Phylogenetic Analysis

#### 3.1.1. Nuclear phylogeny

Our final concatenated nuclear dataset contained 3307 bp from nine nuclear genes: ABCA1 (324 bp), BCHE (244 bp), BDNF (468bp), DMRT1 (530 bp), ERC2 (333 bp), FAM123B (369 bp), FES (479 bp), LZM (136) and MAPKAP (396 bp) for 20 individuals.

In the Bayesian phylogeny (Fig. 1), the CG lineage was sister to *Trachypithecus* with a strong support (PP = 1). The *Semnopithecus hypoleucos* sequence from this study branched with the rest of the *Semnopithecus* individuals. Furthermore, *Semnopithecus* and *Trachypithecus* were sister to each other to the exclusion of the odd-nosed monkey group. Relationships within these major clades are not completely resolved and lack good support. The ML tree (not shown) also retrieved similar relationships with bootstrap support (BS = 98) for the CG + *Trachypithecus* clade.

#### 3.1.2. Mitochondrial Phylogeny

The mitochondrial dataset consists of a large fragment (5933 bp) containing eight mitochondrial genes (Table 2) from 21 individuals. In the Bayesian tree (Fig. 2), capped langur (from the CG lineage group) was sister to *T*. *shortridgei* and together these two species are sister to other *Trachypithecus* species from Southeast Asia (PP=1). Furthermore, *Presbytis melalophos*, instead of *Semnopithecus*, was sister to *Trachypithecus*. Here we see discordance between mtDNA and nuclear dataset with respect to relationships between *Semnopithecus, Trachypithecus* and *Presbytis* (For further discussion see Roos et al., 2011). The topology in the likelihood tree (BS = 100; not shown) was similar to the Bayesian tree. Thus, both nuclear and mitochondrial markers suggested sister relationship between CG lineage and *Trachypithecus*. We also conducted a separate analysis of the mtDNA dataset that included the previously reported numts by Wang et al. (2015). In this tree, the mtDNA fragment generated in this study did not branch with the numts (tree not shown). Additionally, our sequence did not have any of the typical changes associated with numts, such as indels that cause frame shift, nonsense mutations etc.

### 3.2. Species tree estimation

Our coalescent based multilocus analysis of the nuclear dataset generated contradictory results. In the SVDquartet analysis, where multispecies coalescent model was used without taxon partitions, CG lineage was sister to *Trachypithecus* (BS = 36) with the exclusion of *Semnopithecus*. However, when “taxon partitions” option was used, the position of CG lineage was unresolved with respect to *Trachypithecus* and *Semnopithecus*. Whereas, in the result from the ASTRAL tree, *Semnopithecus* was sister to *Trachypithecus* with the exclusion of CG lineage with low branch support (PP = 0.42). (Figure 3).

**Fig. 3:**
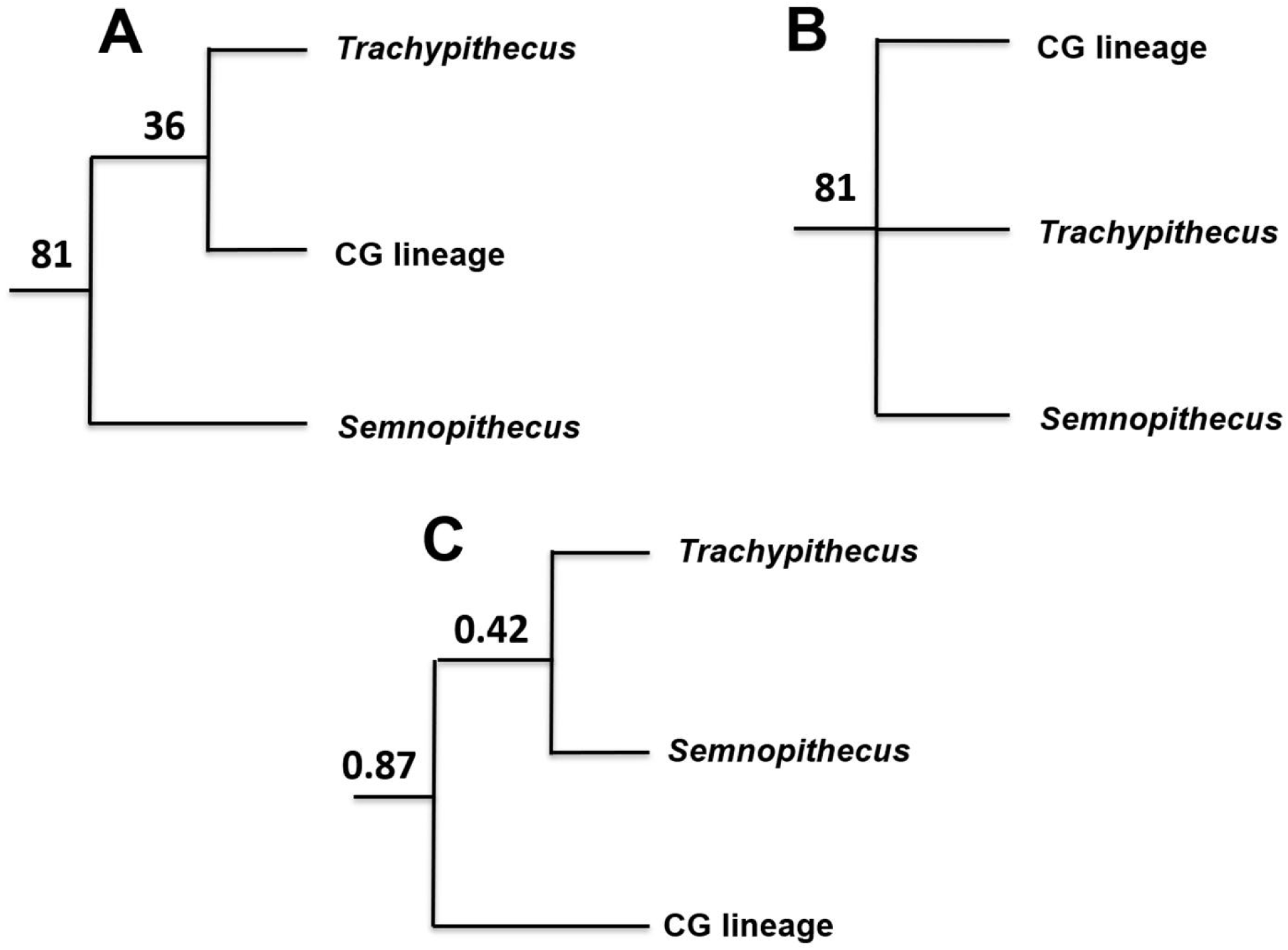
The phylogenetic position of CG lineage with respect to *Semnopithecus* and *Trachypithecus* – from different analyses. (A) species-tree topology from SVDq without taxa partitions (for details, see text), (B) topology estimated with SVDq with taxa partitions (for details, see text), (C) species-tree topology estimated with ASTRAL-II. Numbers at the node are support for that node (support values only for node of interest are shown). Panel A & B shows the BS values; panel C shows PP values.

### 3.3. Divergence Dating

Molecular dates are shown in Fig. 4. These age estimates show the split of African and Asian colobines to be 9.5 mya with CI of 12.6 – 7.09 mya. Within the Asian colobines, the odd-nosed monkeys are estimated to have diverged from the langurs in late Miocene between 9.8 and 6.7 mya. The split between *Semnopithecus* and *Trachypithecus* is estimated to be between 5.2 and 2.7 mya. The date estimates in this study are younger than estimated in an older study using nuclear concatenated dataset (Perelman et al., 2011). Divergence of CG lineage from the rest of *Trachypithecus* occurred 2.5 mya with CI of 4.0 -1.2 mya. The ESS values for all the parameters were >>200 for the two independent BEAST runs suggesting stationarity.

**Fig. 4:**
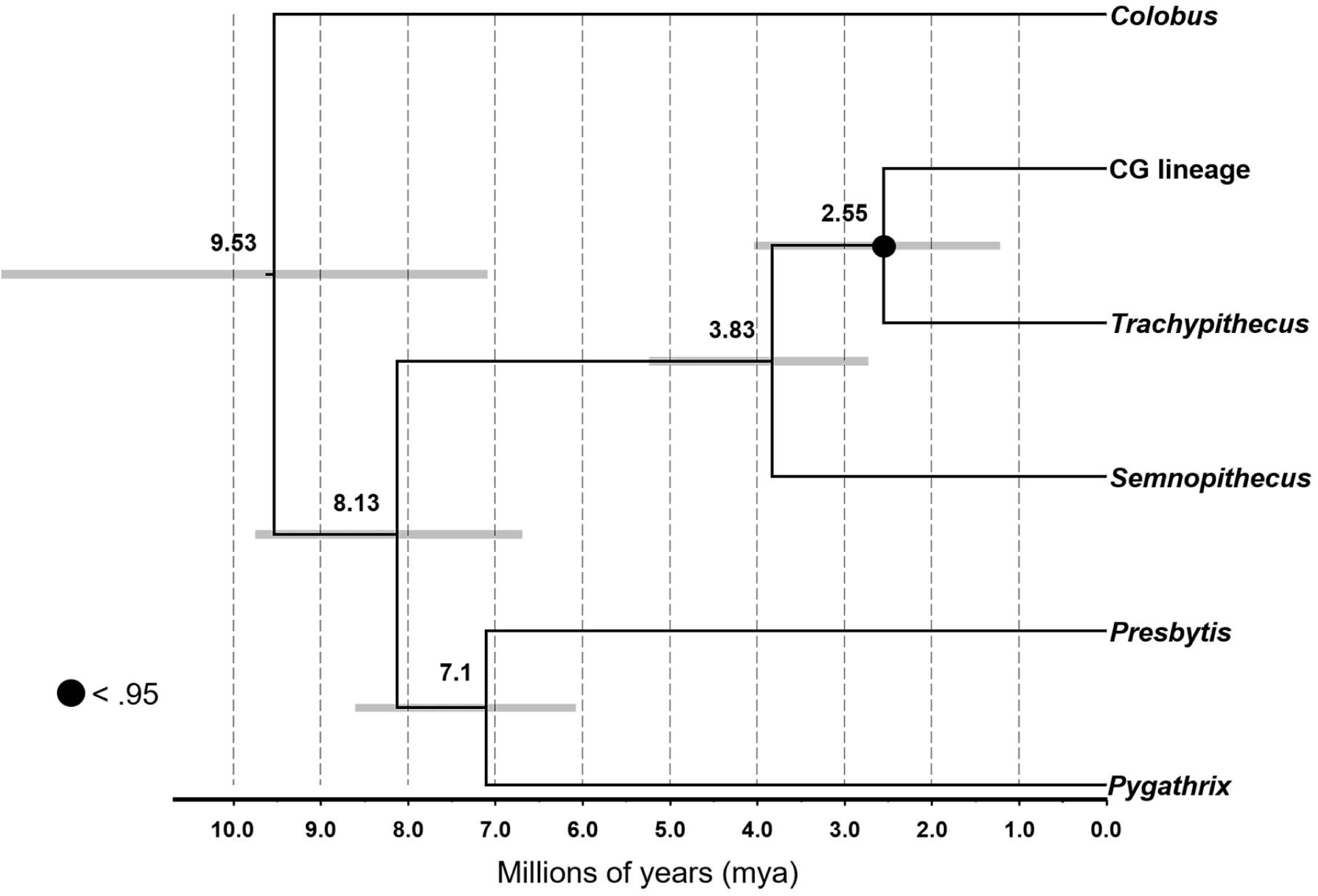
Divergence time tree constructed using coalescent approach in *BEAST using the nuclear data set. The value above each node indicates the mean divergence time and the grey bars indicate the 95% credible intervals. PP support for all the nodes was above 0.95 (except where indicated).

## 4. Discussion

Previous studies that have attempted to resolve the phylogenetic placement of CG lineage were based on single nuclear and / or mitochondrial markers (Karanth et al., 2008; Osterholz et al., 2008; Wangchuk et al., 2008). Among these studies, Karanth et al. (2008) and Osterholz et al. (2008) reported incongruence between nuclear and mitochondrial markers which was taken as evidence for hybrid origin of CG lineage. However, the study by Wang et al. (2015) suggested that this incongruence might be due to the use of numts instead of true mitochondrial sequences. In the present study, we used multiple nuclear and mitochondrial markers to resolve the phylogenetic position of CG lineage. Additionally, we also implemented both concatenated analysis and coalescent based species tree building methods to better understand the origin of CG lineage (Fig. 3).

Phylogenetic tree obtained from the nuclear dataset showed the CG lineage to be monophyletic. However, within CG lineage, neither capped or golden langur was found to be monophyletic. This could be because these are recently diverged taxa and the nuclear DNA does not contain enough information to separate out these two lineages. The relationship within CG lineage can further be resolved by using fast evolving nuclear markers, such as microsatellites. Additionally, adding more samples to the phylogeny might also help resolve the relationship within the CG lineage.

The nuclear as well as the mitochondrial concatenated analyses placed CG lineage sister to *Trachypithecus* with high support. However, the nuclear species trees supported conflicting topologies. Among them only SVDquartet analysis (without taxon partitions) recovered the above relationship but with very low support (Fig 3, A). Whereas ASTRAL and SVDquartet (with taxon partitions) supported alternate scenarios again with low support. Thus, the species trees are unable to resolve the relationship between *Trachypithecus, Semnopithecus* and CG lineage. The low support for relevant nodes in the species trees is suggestive of different evolutionary histories of these nuclear markers due to processes like incomplete lineage sorting. Whereas the high support for CG lineage + *Trachypithecus* monophyly in the nuclear concatenated analysis might be due to few highly variable markers overwhelming the dataset. Additionally, under conditions of low ILS, the species trees are identical to trees based on concatenated analysis (Chou et al., 2015) which is not the case with our results.

Collectively, these results suggest that the phylogenetic position of CG lineage remains unresolved. The convoluted evolutionary history of the CG nuclear makers, we believe, is due to hybridization rather than ILS. This is because, multispecies coalescent models incorporate ILS in species tree estimation (Chou et al., 2015) but these methods cannot distinguish between hybridisation and ILS (Kubatko, 2009). There are three additional lines of evidence that support this scenario. First the numts analyses by Karanth (2008) and Wang et al. (2015) strongly support past hybridization between *Semnopithecus* and *Trachypithecus*. Secondly, the CG lineage is distributed in an area that abuts the eastern and western distributional limits of *Semnopithecus* and *Trachypithecus* respectively. Recent discovery of the late Pliocene fossil of *Semnopithecus gwebinensis* from central Myanmar (Takai et al., 2015) suggests that *Semnopithecus* langurs were distributed till central Myanmar in late Pliocene, whereas today the eastern most distribution of *Semnopithecus* is Bangladesh and Bhutan. This implies that there was an overlap in the distribution of *Semnopithecus* and *Trachypithecus* in the past. Thirdly, the mean body weight (Wang et al., 2015), and body size and skull measurements of CG lineage is intermediate between *Semnopithecus* and *Trachypithecus* (Table S6). Presence of morphologically intermediate forms can be an indication of possible hybridization (Zinner et al., 2011). Such intermediate forms have been seen in other primates like baboons where hybridization has been reported (Phillips-conroy & Jolly, 1981).

Wang et al. (2015) suggest that this hybridization could have been initiated by the bigger *Semnopithecus* males which have an advantage over the smaller *Trachypithecus* males for access to *Trachypithecus* females. The genetic data is consistent with this scenario wherein the mtDNA of CG lineage is related to *Trachypithecus* whereas the nuclear DNA appears to be of both *Semnopithecus* and *Trachypithecus* origin. Our dating analysis (Fig. 4) shows the divergence between *Semnopithecus* and the *Trachypithecus* + CG clade to be around early to mid-Pliocene (3.9 mya; CI – 5.2 mya to 2.7 mya) and the divergence between *Trachypithecus* and CG lineage occurred in late Pliocene (2.6 mya; CI – 4.0 mya to 1.2 mya). Thus, this hybridisation event between lineages of *Semnopithecus* and *Trachypithecus* might have occurred during the Pliocene to early Pleistocene period (5.2 mya to 1.2 mya).

## 5. Conclusion

Concatenation based analysis of nuclear markers yielded tree with good node support for the placement of CG lineage sister to *Trachypithecus*. However, results from different MSC method based analyses were incongruent with each other and with the concatenated tree suggesting ILS/ hybridization in the data set. The low node support in the trees for the node of interest (Fig. 3) could also arise due to gene tree estimation error owing to splitting the overall dataset into small partitions (Gatesy & Springer, 2013; Meredith et al., 2011; Simmons & Gatesy, 2015). Further analysis using longer gene sequences to build individual gene trees, can help resolve this issue. Nevertheless, we suggest the use of both the concatenation and coalescent methods for a robust understanding of various questions in evolutionary biology and systematics.

Overall results from this study and previous studies, suggests that CG lineage might have evolved as a result of hybridisation between *Semnopithecus* and *Trachypithecus*. The genetic evidence for the reticulate evolution of CG lineage needs to be further fortified with more nuclear markers. Nevertheless, the discordance between concatenated and species tree approaches with respect to the position of CG lineage implies that this taxon has a unique evolutionary history. Currently, the CG lineage is placed in the genus *Trachypithecus*, but more studies need to be undertaken to resolve its taxonomic position.

## Supporting information

Supplementary information.docx

## Supplementary Information

Supporting data is in the file Supplementary information.docx

## Acknowledgments

We thank the Department of Biotechnology, Govt. of India and the Ministry of Environment Forest and Climate Change for the funds that covered the molecular work and a part of fieldwork for this study. The forest department of Assam provided the permits to collect samples. We thank Guwahati zoo for allowing us to collect the samples. Dr. Senthil Kumar and Dr. H.T. Lalremsanga from Mizoram University for providing samples from Tripura zoo and Mizoram zoo. Dr. G. Umapathy from LaCONES, CCMB for help in sample collection from Hyderabad zoo. KA would like to thank Mr. Rajani Deka and Mr. Firoz Ahmed for support during field work. Aniruddha Datta-Roy, Ishan Agarwal and V. Deepak for comments on the manuscript. Bhavani from CES for help in accounting.

