## Supplementary information.docx for "The convoluted evolutionary history of the capped-golden langur lineage (Cercopithecidae: Colobinae) – concatenation versus coalescent analyses"

**Table S1:** List of specimens used in the mtDNA phylogeny.

| **Name** | **GenBank Accession numbers** |
| --- | --- |
| *Trachypithecus shortridgei* | HQ149048 |
| *Trachypithecus germaini* | HQ149047 |
| *Trachypithecus hatinhensis* | HQ149046 |
| *Trachypithecus cristatus* | KJ174503 |
| *Trachypithecus obscurus* | AY863425 |
| *Trachypithecus francoisi* | KJ174502 |
| *Trachypithecus shortridgei* (wang2015) | KP834334 |
| *Trachypithecus pileatus* | KF680163 |
| *Semnopithecus entellus* | DQ355297 |
| *Semnopithecus johnii* | HQ149050 |
| *Semnopithecus vetulus* | HQ149049 |
| *Presbytis melalophos* | DQ355299 |
| *Rhinopithecus roxellana* 1 | JQ821835 |
| *Rhinopithecus roxellana* 2 | DQ355300 |
| *Rhinopithecus avunculus* | HM125578 |
| *Pygathrix nigripes* | JQ821840 |
| *Pygathrix nemeaus* | DQ355302 |
| *Nasalis larvatus* | JF293094 |
| *Nasalis concolor* | JF293095 |
| *Colobus guereza* | AY863427 |
| ^†^Capped langur (CES 11/299) | - |

† - Sequenced in this study

**Table S2:** List of specimens used in the nuclear data set.

|  | | Genes | | | | | | | | |
| --- | --- | --- | --- | --- | --- | --- | --- | --- | --- | --- |
| Sr. No. | **Species** | **ABCA1** | **BCHE** | **BDNF** | **DMRT1** | **ERC2** | **FAM123B** | **FES** | **LZM** | **MAPKAP** |
| 1 | *S. e. entellus* | HM765389 | HM764193 | HM763811 | HM762646 | HM762230 | HM762051 | HM761726 | AF294862 | HM760733 |
| 3 ^†^ | *S. hypoleucos* CES08/333 |  |  |  |  |  |  |  |  |  |
| 4 | *S. vetulus* | HM765412 | HM764189 | HM763819 | HM762672 | HM762231 | HM762061 | HM761734 | AH004926 | HM760757 |
| 5 | *T. phayrei* | HM765410 | HM764161 | HM763818 | HM762670 | HM762237 | HM762060 | HM761733 | AF294867 | HM760755 |
| 6 | *T. obscurus* | HM765409 | HM764066 | HM763817 | HM762669 | HM762248 | HM762059 | HM761732 | U76917 | HM760754 |
| 7 | *P.melalophos* | HM765375 | HM764081 | HM763806 | HM762629 | HM762234 | HM762046 | HM761722 | ***** | HM760717 |
| 8 | *R. brelichi* | HM765387 | HM764153 | HM763810 | HM762642 | HM762235 | HM762050 | HM761725 | ***** | HM760729 |
| 9 | *N. larvatus* | HM765362 | HM764154 | HM763800 | HM762616 | HM762243 | HM762040 | ***** | U76945 | HM760705 |
| 10 | *Py.nigripes* | HM765377 | HM764167 | HM763809 | HM762631 | HM762219 | HM762049 | ***** | ***** | HM760719 |
| 11 | *Py. nemaeus* | HM765376 | HM764159 | HM763808 | HM762630 | HM762229 | HM762048 | HM761724 | U76941 | HM760718 |
| 12 | *Py. cinerea* | HM765369 | HM764165 | HM763807 | HM762623 | HM762251 | HM762047 | HM761723 | ***** | HM760711 |
| 13 | *C. guereza* | HM765292 | HM764095 | HM763784 | HM762539 | HM762317 | HM762019 | HM761695 | U76916 | HM760637 |
| 14 ^†^ | *S. hypoleucos* CES09/401 |  |  |  |  |  |  |  |  |  |
| 15 ^†^ | Capped langur CES 12/300 |  |  |  |  |  |  |  |  |  |
| 16 ^†^ | Capped langur CES 11/299 |  |  |  |  |  |  |  |  |  |
| 17 ^†^ | Golden langur CES 12/303 |  |  |  |  |  |  |  |  |  |
| 18 ^†^ | Golden langur CES12/305 |  |  |  |  |  |  |  |  |  |
| 19 ^†^ | Capped langur CES12/308 |  |  |  |  |  |  |  |  |  |
| 20 ^†^ | Capped langur CES12/309 |  |  |  |  |  |  |  |  |  |

† - Sequenced in this study

**Table S3:** Partition schemes and best-fit models of sequence evolution selected in PartitionFinder for reconstructions of phylogeny using Bayesian inference (MrBayes), maximum likelihood (RAxML) and divergence dating in BEAST. Codon positions are denoted by cp1, cp2 and cp3. E1 = Exon 1; E2 = Exon 2; **(A)** = Nuclear markers; **(B)** = Mitochondrial markers.

**(A)**

|  | Partition no. | Partition name | Best substitution model | |
| --- | --- | --- | --- | --- |
|  |  |  | **MrBayes** | **RAxML** |
|  | 1 | ABCA1_intron | HKY | GTR + G |
|  | 2 | BCHE cp1 | HKY + I | GTR + G |
|  | 3 | BCHE cp2 | F81 | GTR + G |
|  | 4 | BCHE cp3 | HKY | GTR + G |
|  | 5 | BDNF cp1 | F81 | GTR + G |
|  | 6 | BDNF cp2 | JC69 | GTR + G |
|  | 7 | BDNF cp3 | HKY + I | GTR + G |
|  | 8 | DMRT1_intron | HKY | GTR + G |
|  | 9 | ERC2_intron | HKY | GTR + G |
|  | 10 | FAM123B cp1 | HKY + I | GTR + G |
|  | 11 | FAM123B cp2 | HKY + I | GTR + G |
|  | 12 | FAM123B cp3 | HKY + I | GTR + G |
|  | 13 | FESE1 cp1 | HKY + I | GTR + G |
|  | 14 | FESE1 cp2 | F81 | GTR + G |
|  | 15 | FESE1 cp3 | HKY + I | GTR + G |
|  | 16 | FES_intron | HKY + I | GTR + G |
|  | 17 | FESE2 cp1 | HKY + I | GTR + G |
|  | 18 | FESE2 cp2 | JC69 | GTR + G |
|  | 19 | FESE2 cp3 | HKY | GTR + G |
|  | 20 | LZM | HKY + I | GTR + G |
|  | 21 | MAPKAP_intron | HKY | GTR + G |
| BEAST | 1 | ABCA1 | HKY+ I | |
|  | 2 | BCHE | HKY+ I | |
|  | 3 | BDNF | HKY | |
|  | 4 | DMRT1 | HKY+ I | |
|  | 5 | ERC2 | HKY+ I | |
|  | 6 | FAM123B | HKY + I | |
|  | 7 | FES | HKY | |
|  | 8 | LZM | HKY | |
|  | 9 | MAPKAP | HKY+ I | |

**(B)**

| Partition no. | Partition name | Best substitution model | |
| --- | --- | --- | --- |
|  |  | **MrBayes** | **RAxML** |
| 1 | COX1_cp1 | HKY+I+G | GTR+I+G |
| 2 | COX1_cp2 | HKY+I+G | GTR+I+G |
| 3 | COX1_cp3 | HKY+G | GTR+I+G |
| 4 | COX2_cp1 | HKY+G | GTR+I+G |
| 5 | COX2_cp2 | HKY+G | GTR+I+G |
| 6 | COX2_cp3 | HKY+G | GTR+I+G |
| 7 | ATP8_cp1 | HKY+I+G | GTR+I+G |
| 8 | ATP8_cp2 | HKY+G | GTR+I+G |
| 9 | ATP8_cp3 | HKY+I+G | GTR+I+G |
| 10 | ATP6_cp1 | HKY+G | GTR+I+G |
| 11 | ATP6_cp2 | HKY+I+G | GTR+I+G |
| 12 | ATP6_cp3 | GTR+I | GTR+I+G |
| 13 | COX3_cp1 | HKY+I+G | GTR+I+G |
| 14 | COX3_cp2 | HKY+I+G | GTR+I+G |
| 15 | COX3_cp3 | HKY+G | GTR+I+G |
| 16 | ND3_cp1 | HKY+G | GTR+I+G |
| 17 | ND3_cp2 | HKY+I+G | GTR+I+G |
| 18 | ND3_cp3 | GTR+I | GTR+I+G |
| 19 | ND4L_cp1 | HKY+G | GTR+I+G |
| 20 | ND4L_cp2 | HKY+I+G | GTR+I+G |
| 21 | ND4L_cp3 | GTR+I | GTR+I+G |
| 22 | ND4_cp1 | HKY+I+G | GTR+I+G |
| 23 | ND4_cp2 | HKY+G | GTR+I+G |
| 24 | ND4_cp3 | HKY+I+G | GTR+I+G |

**Table S4:** Samples used in this study.

| **Sample ID** | **Species** | **Sample collected from** | **Location in wild** | **Sample Type** |
| --- | --- | --- | --- | --- |
| CES11/299 | Capped langur | Hyderabad Zoo | Nagaland | Tissue |
| CES12/300 | Capped langur | Hyderabad zoo | Unknown | Tissue |
| CES12/303 | Golden langur | Guwahati Zoo | Kachugaon, Assam | Fecal |
| CES12/305 | Golden langur | Guwahati Zoo | Bongaigaon, Assam | Fecal |
| CES12/308 | Capped langur | Guwahati Zoo | Guwahati, Assam | Fecal |
| CES12/309 | Capped langur | Guwahati Zoo | Amchang, Assam | Fecal |

**Table S5:** *A priori* species assignment in the respective genera for the dating analysis using starBEAST in BEAST v2.4.7.

| ***Lineage** | **Species** |
| --- | --- |
| CG lineage | CL_CES11/299 |
|  | GL_CES12/303 |
| *Colobus* | *Colobus polykomos* |
|  | *Colobus guereza* |
| *Presbytis* | *Presbytis comata* |
|  | *Presbytis melalophos* |
| *Pygathrix* | *Pygathrix cinerea* |
|  | *Pygathrix nemaeus* |
| *Semnopithecus* | *Semnopithecus entellus* |
|  | *Semnopithecus vetulus* |
| *Trachypithecus* | *Trachypithecus obscurus* |
|  | *Trachypithecus phayrei* |

**Table S6:** Skull measurements and body size measurements (Khajuria, 1956; Pocock, 1939).

| Genus | Species | Skull measurements (mm) | | | Body length (cm) | Tail length (cm) |
| --- | --- | --- | --- | --- | --- | --- |
|  |  | **Total length** | **Condylobasal length** | **Zygomatic width** |  |  |
| *Semnopithecus* | *Semnopithecus schistaceus* | 141 | 113 | 108 | 69.8 | 99 |
|  | *Semnopithecus ajax* | 144 | 113 | 110 | 76.2 | 96.5 |
|  | *Semnopithecus entellus* | 130 | 106 | 106 | 64.7 | 107.9 |
|  | *Semnopithecus hypoleucos* | 120 | 91 | 92 | 68.5 | 109.2 |
|  | *Semnopithecus priam* | 123 | 99 | 96 | 64.2 | 95.2 |
| CG lineage | Capped langur | 116 | 92 | 88 | 69.5 | 99 |
|  | Golden langur | 110.5 | 88.1 | 89.1 | 72 | 90 |
|  | Shortridge’s langur | 113 | 91 | 91 | 71.6 | 103.6 |
| *Trachypithecus* | *Trachypithecus phayrei* | 106 | 87 | 80 | 57.1 | 78.7 |
|  | *Trachypithecus obscurus* | 106 | 84 | 78 | 59.6 | 71.1 |
